# Drug safety in pregnancy: review of study approaches required by regulatory agencies

**DOI:** 10.1101/640128

**Authors:** Andrea V Margulis, Mary Anthony, Elena Rivero-Ferrer

## Abstract

**Purpose of review:** We reviewed postauthorization pregnancy safety studies requested by regulatory agencies to explore which study approaches have been typically requested and to what extent these have changed over time.

**Recent findings:** The most common study approach requested by the US Food and Drug Administration (FDA) is pregnancy exposure registries (observational cohorts with prospective data collection), per the FDA’s Postmarketing Requirements and Commitments (PMR/PMC) database. Since 2017, this requirement has often been paired with a request for a database study (observational study using preexisting electronic health care data), both approaches assessing pregnancy and fetal outcomes. From studies registered in the European Union electronic Register of Post-Authorisation Studies, we observed a similar number of pregnancy exposure registries and database studies, both approaches also assessing pregnancy and fetal outcomes. In requests for drugs approved since 2014, preference appears to have shifted toward studies using preexisting electronic health care databases from multiple countries.

**Summary:** Pregnancy exposure registries have been the most commonly required study approach on drug safety in pregnancy. Recent regulatory requests and activities denote an increasing interest in other approaches.

## INTRODUCTION

Prescription drugs and biologics are approved to go into the market after animal studies and randomized clinical trials have shown efficacy and a favorable benefit-risk profile. Regulatory agencies require that pharmaceutical companies conduct postauthorization safety studies (PASSs) to collect additional information on potential safety issues or subpopulations that were missing or underrepresented during the product development program. An important aspect of these postauthorization studies is that they can be done using real-world data to assess safety of the medical products among patients that might have been excluded in the preauthorization trials.

The FDA may require postauthorization pregnancy registries when it is likely that the product will be used by women who are or may become pregnant or there is a potential safety concern based on the pharmacologic class, data from animal studies or clinical trials, or human case reports [1]. Pregnancy exposure registries are designed and established to prospectively collect detailed information on pregnancy exposures and outcomes (they may also collect some information retrospectively), often using questionnaires and medical record review. Patient enrollment depends on whether patients and health care providers are aware of the existence of the registry and are willing to enroll and provide information. It generally takes many years for pregnancy exposure registries to reach their prespecified study size. Often, pregnancy exposure registries do not include an unexposed population and results (such as the proportion of infants with congenital malformations) are compared with external references.

Retrospective cohorts or case-control studies that use existing health care databases, such as electronic medical records and health care claims, are also commonly used for research on drug safety in pregnancy. In these data sources, information is collected prospectively as patients receive health care services, but studies are designed and conducted after exposure and outcomes have occurred and been recorded. Patient-level information may contain unconfirmed elements and data may be less detailed than that obtained with primary, prospective data collection, but study sizes can be larger and adjusted measures of association can be estimated. Another data source for case-control studies on congenital malformations is case-control surveillance data, which include detailed retrospectively collected information on pregnancy exposures, maternal characteristics and pregnancy outcomes for malformed infants and non-malformed controls.

Other approaches that have been used for postauthorization drug safety research include exposure or disease registries, in which patients who use the product or have the condition for which the product is indicated are enrolled and followed up. These studies are generally able to provide a more complete view of the safety profile of the drug or the burden of illness of the disease but are less detailed with regard to pregnancy exposures, clinical course, and outcomes than other designs. Another approach calls for continuing the follow-up of spontaneous reports; information collected by these means is generally not too detailed, pregnancy outcomes are often reported retrospectively, and loss of follow-up is common. Furthermore, the underlying population of users to estimate rates or proportions is not identifiable with this approach.

Here, we used publicly available data sources to review the pregnancy-related PASSs requested by two regulatory agencies, the FDA and European Medicines Agency (EMA), and to explore which study approaches are typically required and to what extent requirements have changed over time. This information may be useful to the pharmaceutical industry to inform them about potential regulatory requirements for products in the pipeline, to researchers to contextualize any given request, and to the clinical community to educate them about the efforts in place to evaluate the safety of medications in pregnancy.

## METHODS

We searched the FDA’s Postmarketing Requirements and Commitments (PMR/PMC) database and the European Union electronic Register of Post-Authorisation Studies (EU PAS Register) to identify regulatory-required pregnancy safety studies using all the information available to date. We summarized key study characteristics overall and in three 6-or 7-year intervals.

### FDA’s Postmarketing Requirements and Commitments Database

The FDA’s PMR/PMC database collects information on studies that were required by the FDA (postmarketing requirements [PMRs]) or that the manufacturer committed to conduct although they were not required (postmarketing commitments [PMCs]) after marketing authorization of drugs or biological products [2]. These PMR/PMCs can be the result of negotiations between the FDA and the pharmaceutical company and can refer to studies that were ongoing at the time of the request. The database is maintained by the FDA and is publicly available for download (https://www.accessdata.fda.gov/scripts/cder/pmc/index.cfm). We downloaded the database that included data through December 31, 2018, and conducted a search using key words. We identified all PMR/PMC descriptions containing the string “pregn” (case insensitive). When we found more than one entry that seemed to refer to a single request, we retained only one (including PMR/PMCs for different forms for the same product, such as tablets and powder for oral suspension; entries with different wording but that seem to describe the same study; and entries that seemed to have been updated by a subsequent entry). We categorized studies as pregnancy exposure registries, health care database studies (including retrospective cohorts and case-control studies), exposure or disease registries, or spontaneous report follow-up studies. Key data elements were extracted into evidence tables. For requests that referred to an ongoing study, we obtained additional information from ClinicalTrials.gov and from documents from the FDA website (https://www.fda.gov). The time from the NDA approval date to the final report due date was calculated.

### EU PAS Register

The EU PAS Register is the mandatory repository for observational postauthorization studies requested by the EMA; registration became mandatory in 2012. Studies requested by other regulatory agencies and studies conducted voluntarily by pharmaceutical companies can also be registered. The database is hosted by the EMA and the information on PASSs is registered by pharmaceutical companies, contract research organizations, or academic centers conducting the research. Protocols, reports, and publications can be uploaded. The publicly available search interface contains several fields that allow the user to identify studies based on study characteristics that are provided by the registrant during study registration (http://www.encepp.eu/encepp/studySearch.htm). We conducted two searches. In the first search, we used the field “Study requested by a regulator” and selected all countries except the US; from the list in the field “Other population,” we selected “Pregnant women”; and from the field “Scope of the study,” we selected “Disease epidemiology,” “Risk assessment,” and “Drug utilization study.” No other filter was implemented. The search was conducted on March 11, 2019, and retrieved 64 studies. The second search was conducted using the search term “pregnancy” in the free-text field “Medical condition”; selecting the options “Active surveillance,” “Observational study,” and “Other” in the field “Study type”; and ticking the field “Study requested by a regulator.” This search was conducted on April 15, 2019, and retrieved 17 studies. We reviewed all records, removed duplicates (study entries identified in both searches), and identified entries for studies that are not part of a risk management plan or that were conducted to satisfy a PMR/PMC for the FDA only. Data were extracted as described in the previous paragraph. Dates of marketing authorization were obtained from the EMA website (https://www.ema.europa.eu/en/medicines).

## RESULTS

### FDA’s Postmarketing Requirements and Commitments Database

Of the 2,663 entries in the PMR/PMC database, our search identified 112 records with the string “pregn”; we removed eight duplicates, six animal studies and two requests for randomized clinical trials. The final dataset consisted of 96 records for 83 products with NDA approval dates between August 1998 and August 2018 (Table 1 and Supplementary material). The number of pregnancy-related PMR/PMCs per year increased over time (Fig 1). Of the 96 records, 57 were for pregnancy exposure registries (average time between NDA approval and final report due date, 11.1 years), 15 were for database studies (9.0 years), 1 was for a study with both components (21.1 years), 15 were for exposure or disease registries (13.8 years), and 3 were for spontaneous report follow-up studies (9.1 years). Planned study duration shortened over time, overall (mean of 16.8 years for drugs approved in 1998-2004 to 9.6 years for drugs approved in 2012-2018) and for each specified study approach. Of the 15 exposure and disease registries, 7 were for products approved in the midthird of the study observation period and 6 were for products approved in the last third. Twelve of 15 database studies were for drugs approved in the last third of the observation period. Twenty records specified that the study was to be conducted in the US, 7 would be international studies, and the remaining 69 records did not specify any geographical focus. Descriptions of two pregnancy exposure registries specified that paternal exposure would be studied.

**Table 1.**
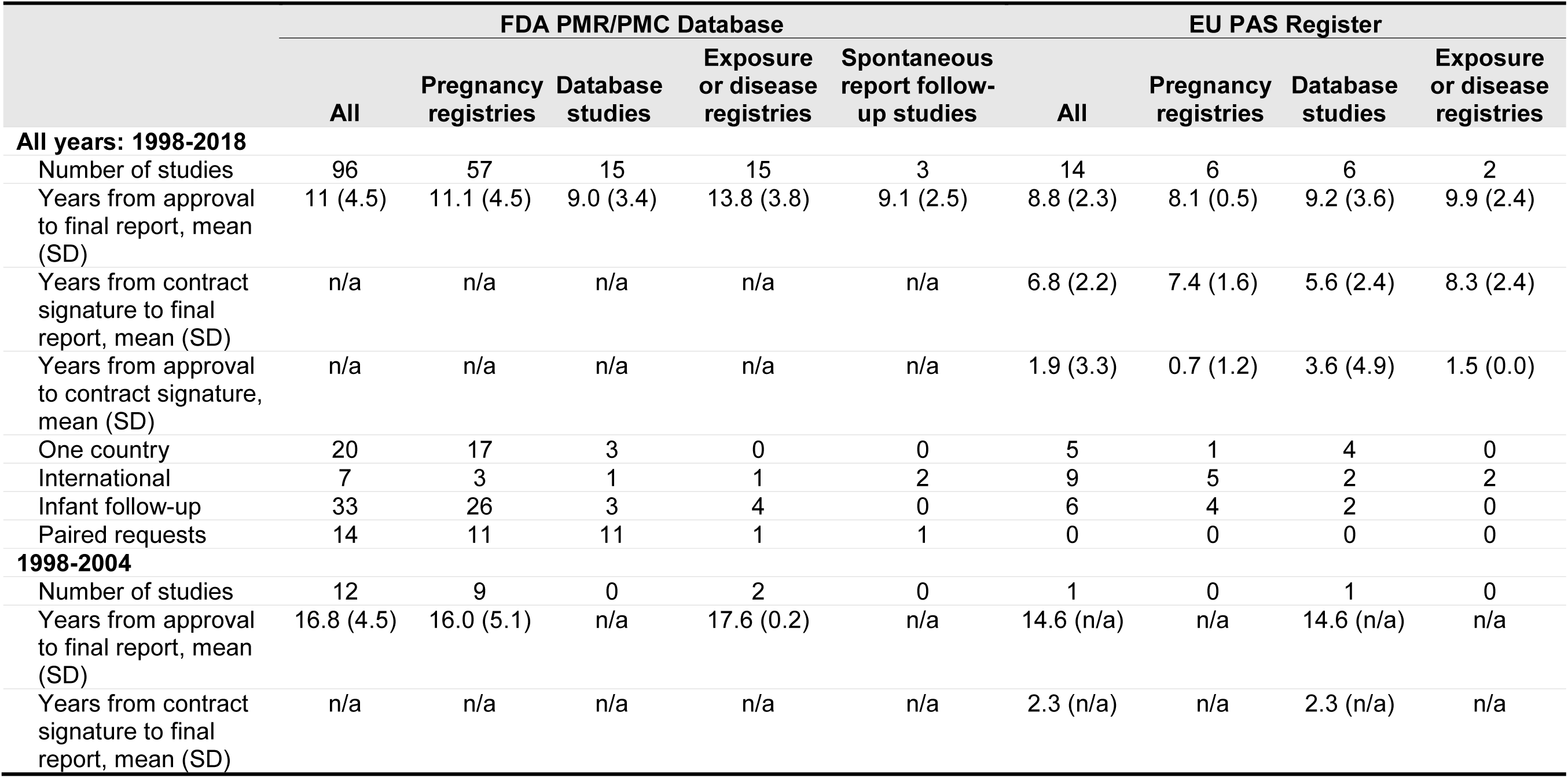
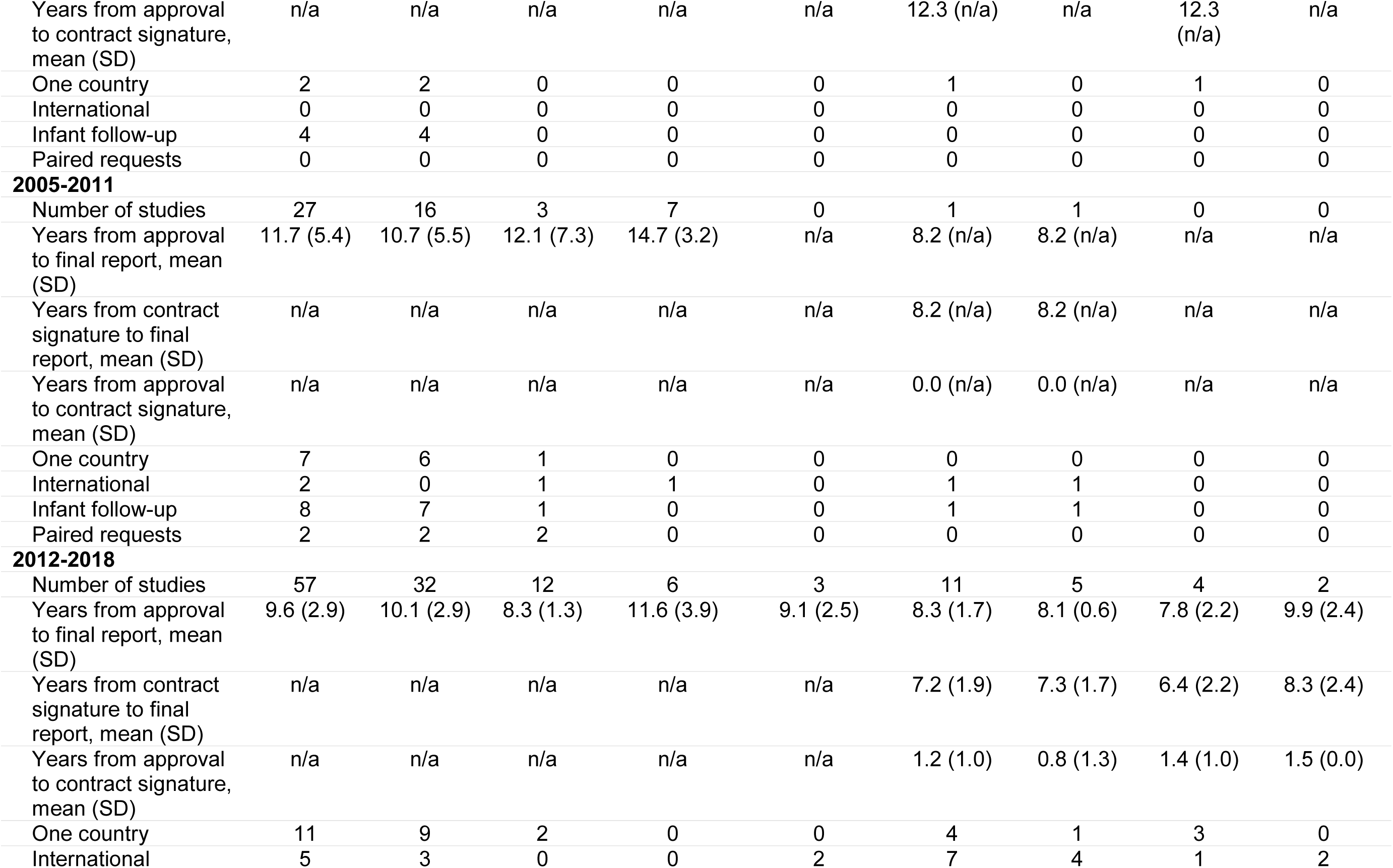
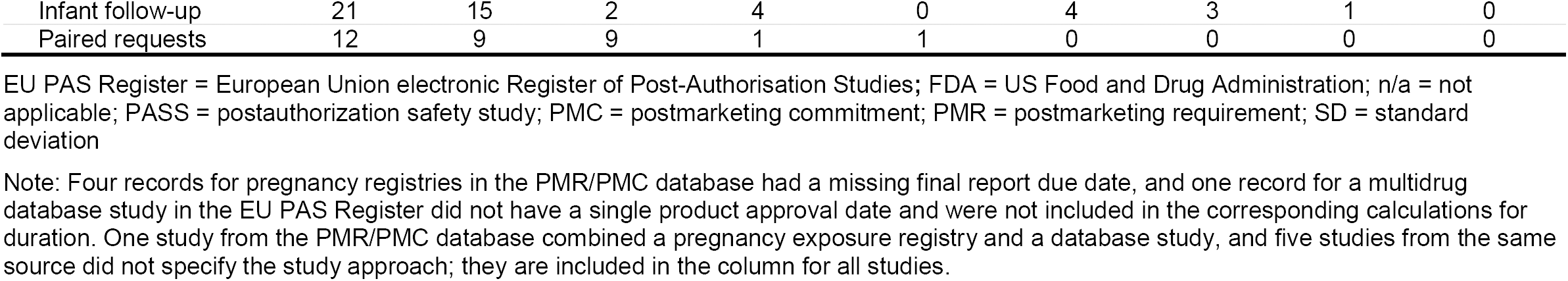
Characteristics of pregnancy PASSs requested by regulatory agencies identified in the FDA PMR/PMC database and the EU PAS Register

**Fig 1.**
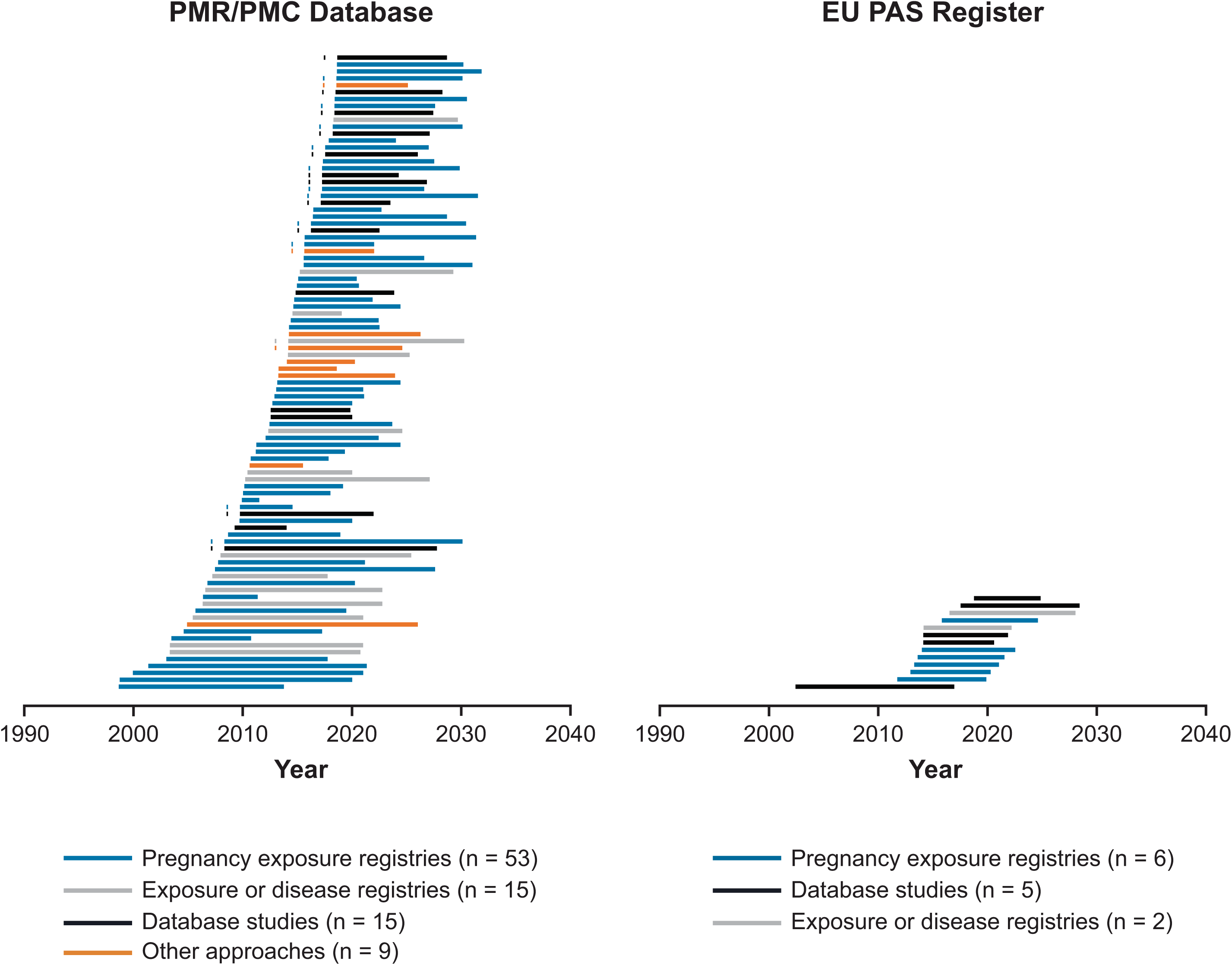
Pregnancy PASSs requested by regulatory agencies identified in the FDA PMR/PMC database and the EU PAS Register: time from drug approval to final report. EU PAS Register = European Union electronic Register of Post-Authorisation Studies; FDA = US Food and Drug Administration; PASS = postauthorization safety study; PMC = postmarketing commitment; PMR = postmarketing requirement. Note: Each segment represents one study. The length of each segment represents the planned study duration, from the new drug application or product approval date to the final report due date. Four records for pregnancy registries in the PMR/PMC database had a missing final report due date, and one record for a multidrug database study in the EU PAS register did not have a single product approval date and were not plotted. Dots to the left of a segment in the left panel indicate that the study is one element of a paired request for a pregnancy exposure registry and a second study with a different design. Also in the left panel, a pregnancy exposure registry for a flu vaccine has a planned duration of 1.6 years

The level of detail in CDER-issued PMR/PMC descriptions was variable, from the request to assess “pregnancy outcomes” in the oldest record to the request “to assess major congenital malformations, spontaneous abortions, stillbirths, preterm births, and small-for-gestational-age births” for the most recent one (this was a very common request in the last 10 years). The Center for Biologics Evaluation and Research (CBER) issued 17 of the 96 PMR/PMCs, mostly for registries.

Descriptions of the five PMR/PMCs for pregnancy exposure registries for multiple-sclerosis (MS) treatments were similar to each other. Descriptions for the two entries for antiepileptic drugs (AEDs) were identical, as were the descriptions for two monoclonal antibodies to treat hypercholesterolemia. Some descriptions were adapted to specific products; for example, for some drugs that can increase the risk of infections in the user, study outcomes included infections in in-utero–exposed infants.

For 11 products, the PMR/PMC database included requests for two studies with different approaches: a pregnancy exposure registry and a second study with a different design. For two additional products (two AEDs), the FDA only required applicants to conduct a study with a design different from that of the North American AED Pregnancy Registry, possibly because the FDA already receives regular reports from that registry. For one additional product, two studies with different approaches were requested, but none was a pregnancy exposure registry. Twelve of these 14 PMR/PMCs pairs were for products approved in 2014-2018. In 7 of the 9 pairs in which the approaches requested were a pregnancy exposure registry and a database study, the time between the NDA approval and final report due date was shorter for the database study (average difference for the 9 pairs, 2.3 years).

### EU PAS Register

Of the 75 unique PASSs identified in the EU PAS Register, 53 were excluded because they did not focus on pregnancy safety and 5 were excluded because they had been requested solely by the FDA (the PMR/PMC database did not include records for 1 of these products when we downloaded it, or currently [search conducted on May 11, 2019]; we identified and included in this review PMR/PMCs for the other products). Three additional studies funded by regulatory agencies and aimed at pregnancy safety were also excluded because they were not required by a risk management plan (two meta-analyses and a database study). Overall, 14 PASSs required by a risk management plan for products approved between 2002 and 2018 were reviewed (Table 1 and Supplementary material); 12 were category 3 (required) and 2 were category 1 (imposed as a condition of marketing authorization). Requestors were the EMA (n = 11) and the EMA and FDA (n = 3). The status of the PASS was finalized for 4 PASSs, ongoing for 8, and planned for 2. Of the 14 PASSs, 6 were pregnancy exposure registries, 6 were cohort studies using automated health care databases, and 2 were exposure or disease registries. Nine studies were international, involving 2 to 20 countries. The PASSs targeted pregnancy outcomes (14 studies), major congenital malformations (13 studies), and maternal outcomes (7 studies). Infants were followed during the 12 months after delivery in 6 studies, and paternal exposure was evaluated in 2 studies.

The time from marketing authorization of the medication in the European Union to the final study report (planned date for ongoing and planned PASSs and for PASSs with early finalization due to drug market withdrawal, and actual date for other finalized PASSs) was, on average, 8.8 years (9.2 for PASSs conducted on health care databases, 8.1 for pregnancy exposure registries, and 9.9 for exposure or disease registries). On average, the duration of the PASS from the date when the funding contract for the study was signed to the date of final study report was 6.8 years, with 5.6 years for PASSs conducted in health care databases, 7.4 years for pregnancy exposure registries, and 8.3 years for exposure or disease registries.

Of the eight entries for drugs approved in 2014 and onward, four were for database studies and two were for exposure or disease registries. Most pregnancy exposure registries were for drugs with older approval dates.

We did not find entries for paired requests for studies with different designs; the two entries for a single drug were for database studies in different locations.

### Studies in Both Sources

The EU PAS Registry listed three studies that had been requested by both the EMA and FDA. For one, a pregnancy exposure registry, we identified and included in this review the corresponding entry in the PMR/PMC database. For another, a paired request (pregnancy exposure registry and a study with a second approach) was added to the PMR/PMC database after the date of our data download. For the third, we could not identify the corresponding entry in the PMR/PMC database (in the version we downloaded or currently [search conducted on May 11, 2019]).

We also identified, for another drug, records in both data sources that appear to refer to the same pregnancy exposure registry. In addition, we identified records for one FDA-requested pregnancy exposure registry for one indication of a drug (study duration, 20 years) and one EMA-requested pregnancy exposure registry for another indication (8 years); both studies are ongoing.

## DISCUSSION

### Main Findings

In the FDA’s PMR/PMC database, we observed an increasing number of requests for pregnancy PASSs. Pregnancy exposure registries were the most commonly requested approach; paired requests for a pregnancy exposure registry and a database study are increasingly common in the US. Among paired requests, the time since NDA approval and the report due date was generally somewhat shorter for the database study. PMR/PMCs in the last 10 years often include a list of key study outcomes, typically major congenital malformations, spontaneous abortions, stillbirths, preterm births, and small-for-gestational-age infants. In the EU PAS Register, we observed a similar number of pregnancy exposure registries and database studies. Requests typically targeted maternal and pregnancy outcomes and congenital malformations. In requests for drugs approved since 2014, preference appears to have shifted toward studies using preexisting electronic health care databases from multiple countries.

### FDA and EMA’s Assessment and Support of Studies With Various Approaches

In 2015, FDA researchers reviewed the planned characteristics of 35 pregnancy exposure registries; the authors stated that well designed pregnancy exposure registries with a sufficiently long enrollment period are valuable, and larger study sizes than typically achieved by pregnancy exposure registries are needed to assess specific major congenital malformations [3]. A follow-up study on 34 of those 35 registries reported a median actual enrollment of 36 pregnancies after a median of 6 years since registry launch, while the median number of worldwide spontaneous reports for use in pregnancy of the medications included in those registries was 450 (15 of the 34 registries were multinational) [4]. The inclusion of the heading “Pregnancy Exposure Registry” at the top of the pregnancy section (Section 8.1) in drug labels confirms the FDA’s continued interest in and support of pregnancy exposures registries [5]. A study co-authored by EMA staff found that of the 62 product pregnancy registries that were listed in risk management plans for centrally authorized products in Europe, 38 were considered uninformative because of low enrollment, or because no protocol or report had been submitted to EMA; the remaining 24 had substantial loss to follow-up or had implemented changes in the enrollment strategy that rendered results uninterpretable, despite the availability of exposed pregnancies as evidenced in spontaneous reports. The authors concluded that these registries, as they are, are not delivering what they are aimed to achieve [6].

The use of existing data sources for drug safety research and, specifically, for research on drug safety in pregnancy, is growing. The FDA has been increasingly interested in incorporating real-world evidence into their decision-making processes, including those related to drug safety in pregnancy [7]. The FDA funded MEPREP, a distributed network of mother-offspring–linked data from the US that included births from 2001-2007 [8], and is developing the Sentinel Pregnancy Tool to enable surveillance of drug use in pregnancy in the Sentinel Distributed Database [9]. The 2019 FDA’s draft Guidance on Postapproval Pregnancy Safety Studies [10], open for public comment at the time of drafting this manuscript, lists database studies as studies that can complement pregnancy exposure registries, confirming that the trend we observed in the US reflects FDA’s current thinking. The EMA commissioned a review of existing data sources for research on drug safety in pregnancy that could be an alternative source to pregnancy registries. This report identified and described a large number of data sources being used for drug safety in pregnancy research [11, 12]. This report was later updated by the EUROmediSAFE Inventory, with 511 entries including pregnancy exposure registries, health care databases and other data sources [13]. In 2017, the EMA issued a call for research organizations to submit proposals for efficacy and drug safety studies using real-world data, for which the focus explicitly listed pregnancy and breastfeeding research [14]. Many other regulatory-funded initiatives that focus or address drug safety in pregnancy and use real-world data exist.

### Results From Pregnancy Exposure Registries and Database Studies

To our knowledge, few studies have specifically compared pregnancy exposure registries and database studies for a given association. In an exercise comparing an international prospective pregnancy exposure registry with a retrospective cohort study based on US claims for years 1996-2012, the database study identified 4,519 sumatriptan-exposed pregnancies, and the registry, 617 [15]. The number of sumatriptan-exposed pregnancies would have been observed in years 2001-2005 in the database study. Among pregnancies with first-trimester sumatriptan exposure, the database study identified a larger proportion of spontaneous and elective abortions than the registry (18% and 5%, respectively, in the database study, and 7% and 3%, respectively, in the registry). This finding illustrates that left truncation can affect pregnancy registries to a larger extent than database studies, as early pregnancy losses sometimes occur before registry enrollment would take place. The proportion of infants with birth defects was identical, 4.0%, in both studies. The pregnancy registry was able to obtain details on dose, indication, and specific congenital malformations. Although this pregnancy registry was conducted to meet a requirement from the FDA, we did not find records for it in the PMR/PMC database (search conducted on May 11, 2019).

A literature review on treatments for MS found that data collected with various approaches was incomplete and lacked standardization [16], and authors recommended the development of a North American MS pregnancy registry to study pregnancy outcomes related to MS and its treatments. This data-quality discussion may apply to studies on other diseases, and the authors cite examples of successful disease pregnancy registries for other conditions. An unrelated research group advocated for a disease registry as the best-positioned design to collect information on the preconceptional course and severity of underlying chronic inflammatory diseases for studies on drug safety in pregnancy [17]. On the other hand, another group of researchers expressed the need for an international multiproduct pregnancy exposure registry for monoclonal antibodies for any indication [18]; this registry would include some but not all treatments for MS. Our group conducted a literature review that identified studies on the safety of MS treatments in pregnancy and identified 43 publications from product-specific pregnancy exposure registries, disease registries, database studies and studies with other approaches [19]. Most studies, regardless of their design, were considered noninformative due to their small study size. Exploration of existing data sources for database studies showed that some existing databases should be capable of supporting informative studies in terms of study size, as long the medication under study has a good uptake [19].

Advantages of paired requests are that the two types of data sources contribute complementary data [20, 10], with pregnancy exposure registries being able to provide detail on exposure and outcomes and database studies being able to obtain narrower safety intervals [21] and/or earlier results. Consistent results from two designs would support the robustness of the findings. Drawbacks include the high cost of the double request, especially when the product is not widely used, creating the need to extend pregnancy exposure registries for long periods, to implement multidatabase studies (as opposed to single database studies), or to conduct international studies. There is a concern that some pregnancies may contribute information to both studies [10], which would be unknown to researchers given current data access and sharing restrictions.

### Limitations and challenges

Limitations of this review include the fact that the two data sources that we used contain different types of information: the FDA’s contains PMR/PMCs, while the EU PAS Register contains information on studies that are being implemented. The delay between the PMR/PMC issuance and the planning and then registering a study may partially explain the smaller number of pregnancy PASS entries in the EU PAS Register. Furthermore, after regulatory requests are issued, pharmaceutical companies may negotiate with regulatory agencies modifications of the study approach based on a study feasibility evaluation. The PMR/PMC description is sometimes succinct, and we applied our interpretation of what was meant to classify the request into one or another approach; readers can see the full PMR/PMC descriptions and our categorization in the Supplementary material. Because of the different structure of both data sources, our searches were organized differently and were not expected to behave identically. Additional data sources to search for safety studies exist, such as ClinicalTrials.gov (https://clinicaltrials.gov/) and the FDA’s Office of Women’s Health list of pregnancy exposure registries (111 entries on May 11, 2019; https://www.fda.gov/scienceresearch/specialtopics/womenshealthresearch/ucm134848.htm). We did not include them in this review because recent studies have looked into them extensively and because registering observational studies in them is not mandatory. Specific aspects of designs may be driven by the indication for each newly approved product; e.g., rare diseases or a crowded therapeutic area with high market fragmentation may determine longer planned study durations. Thus, some trends we observed in this review may reflect trends in NDA filings rather than trends in agencies’ preferences. We found that the PMR/PMC database does not include all studies that would be of interest for this review.

## Conclusions

Pregnancy exposure registries have been the most commonly required study approach on drug safety in pregnancy. Recent regulatory requests and activities denote an increasing interest in other study approaches.

## Acknowledgements

Editorial services were provided by John Forbes and graphic design support was provided by Jason Mathes and Christopher Lovett, all employees of RTI Health Solutions. The authors thank Abenah Harding and Catherine Saltus, also from RTI Health Solutions, for their help in the preparation of this manuscript.

## Human and animal rights

This article does not contain any studies with human or animal subjects performed by any of the authors.

